# Automation of high-throughput workflow for arrayed CRISPR activation library screening

**DOI:** 10.1101/2025.11.10.687722

**Authors:** Chih-Cheng Yang, Aniruddha J. Deshpande, Michael Jackson, Peter D. Adams, Elena B. Pasquale, Rabi Murad, Jiang-An Yin, Yancheng Wu, Anna Beketova, Chun-Teng Huang

**Author notes:** Correspondence – Chun-Teng Huang, Functional Genomics Shared Resources, Sanford Burnham Prebys Medical Discovery Institute, 10901 N. Torrey Pines Road, La Jolla, CA, 92037, T-858-795-5353, E –. Equal contribution (Chih-Cheng Yang, Chun-Teng Huang).

## Abstract

CRISPR-mediated gene activation (CRISPRa) is among the most efficient and reliable strategies for mimicking sustained activation of endogenous promoters and their corresponding genes at physiological levels. By leveraging guide-RNA (gRNA) library design, CRISPRa screens can be applied on a whole-genome scale and are compatible with both arrayed and pooled formats, depending on assay requirements. Compared with conventional arrayed CRISPRa libraries that use single or dual gRNAs and often require multiple gRNA candidates per target, a recently developed CRISPRa library (termed T. gonfio) incorporates four tandem gRNAs per lentivector per target, thereby reducing library complexity and representing the smallest arrayed genome-wide CRISPRa library. To streamline genome-wide arrayed CRISPRa screening, this study developed a high-throughput automated workflow using the Biomek i7 Hybrid liquid-handling platform, integrated with multiple peripheral instruments. The workflow comprises three pipelines: lentiviral library transduction, cell library passaging, and assay processing. These pipelines together establish and maintain the transduced cell library for extended screening times. This enables assay processing at desired extended time points and improves the likelihood of identifying phenotypes that require longer time to develop, making the workflow suitable even for rapidly proliferating cell models. In a pilot arrayed screen using a T. gonfio mini-library targeting kinases and phosphatases, activation of the EPHA2 receptor promoter induced a growth reduction phenotype in the HEK293 cell model. This phenotype was recapitulated in a parallel pooled CRISPRa screen using the same mini-library and further validated in a co-culture assay.

## Introduction

Repurposing CRISPR-Cas9 for transcription regulation by tackling promoter or enhancer activities allows target genes or long non-coding RNAs to be activated via CRISPRa- dCas9-VP64 systems, including SAM [1], suntag [2], split fluorescent protein [3], and others. Commonly, a lentiviral arrayed CRISPR guide-RNA (gRNA) library uses single or dual gRNAs per vector [4]. This system uses the VP64 synthetic transcriptional activator, which includes four copies of the herpes simplex virus VP16 protein domain fused to a deactivated Cas9 protein [3]. It increases the expression of a target gene without altering its coding sequence. Ensuring optimal activation efficiency often requires multiple gRNAs targeting different regions of a promoter, thereby increasing library complexity and adding burden to the screening process. To address this, Yin *et al.* created a human genome- wide arrayed CRISPR activation quadruple gRNAs (qgRNAs) library named T. gonfio library [5]. The size of this library was minimized by designing four tandem Cas9 gRNAs per lentivector, meaning that each lentivirus carries four tandem gRNAs targeting each gene promoter. This library was originally designed for CRISPR gene activation but is also suitable for epigenetic silencing [5].

Arrayed CRISPR library screening offers several compelling benefits, such as broader assay compatibility, higher sensitivity in phenotypic screens, straightforward genotype- phenotype correlation, and a reduced need for data deconvolution [6–9]. However, few major obstacles have prevented it from being widely adopted in research. A typical arrayed CRISPR screen requires at least 7 to 14 days post-transduction before assay readout. This prolonged cell growth time can lead to over-confluent cultures and thus compromise screening results. In addition, successful arrayed library screening relies on the production of high titer lentiviral CRISPR gRNA libraries to ensure efficient transduction in difficult-to-transduce cell types. Therefore, two major hurdles are: 1) the labor-intensive, time-consuming, and costly workflows associated with arrayed mammalian cell culture passaging, and 2) high titer lentiviral CRISPR gRNA library production at a genome-wide scale. To tackle these challenges and streamline the workflow through automation, two previous studies established high-throughput arrayed cell culture passaging and lentivirus production workflows using the Biomek i7 Hybrid automated workstation [10, 11]. Building upon these prior methodologies, in this study we have developed an automated workflow optimized for arrayed CRISPR screening and suitable for both small and large-scale libraries.

The Biomek automation platform demonstrates substantial versatility, as evidenced by its broad utility across a wide range of workflows, including mammalian and stem cell culture at various throughputs [10, 12, 13, 14, 15, 16, 17], high throughput screening [14, 15, 16, 18, 19], assay processing [14, 20, 21, 22], sample treatment and reformatting [23, 24, 25, 26, 27], PCR and qPCR [17, 28, 29, 30, 31], mRNA and ChIP-Seq library preparation [32, 33, 34], genomic and proteomic workflows [35, 36, 37, 38, 39, 40], as well as lentivirus production and titration [11]. The present study implements the Biomek i7 Hybrid automated workstation and utilizes the Biomek 5 software to program and execute sophisticated liquid handling methods. SAMI EX software was integrated on this foundation to simulate, schedule, and coordinate liquid handling with multiple peripheral instruments. This synergistic platform supported three automated pipelines, including lentiviral library transduction, cell library passaging, and assay processing, collectively constituting a high throughput automated workflow optimized for arrayed CRISPRa library screening. The lentiviral library transduction pipeline initiates the screening by generating an arrayed transduced cell library. The second pipeline for cell library passaging allows prolonged extension of the screening by maintaining the cell libraries until the desired phenotype emerges. The third pipeline processes the viability assay and acquires the live-cell data for cell number measurement and normalization. As a proof of concept, we first evaluated CRISPRa efficiency using eight kinase- or phosphatase-targeting qgRNAs derived from the T. gonfio kinase, phosphatase, and drug-target sub-library. We selected genes whose expression levels are relatively low in HEK293 cells compared with other cell lines. After confirming the robustness of CRISPRa, we screened an arrayed mini- library targeting eleven kinases or phosphatases from the T. gonfio sub-library. These genes were selected for their reported growth-related phenotypes upon activation. After identifying the EPHA2 receptor tyrosine kinase as a hit, a co-culture assay and a deconvolution screen were performed to validate the EPHA2-associated growth inhibition phenotype.

## Materials and Methods

### 1. System setup of the Biomek i7 Hybrid automated workstation and integrated instruments

The Biomek i7 automated liquid handling system (Product No. B87585, Beckman Coulter) is equipped with hybrid multichannel pods, a dual-pod system combining 96/384-channel heads and eight independent pipette heads, along with a barcode reader, two track grippers, a plate shuttle station, a Biotek 405 LSHTV Washer (Agilent), a BioShake 3000- T ELM plate shaker (QInstruments), a Cytomat 2C incubator (Thermo Fisher Scientific), a CloneSelect imager (Molecular Devices), and a microcentrifuge (Agilent). These instruments together can automate arrayed CRISPRa screening at high throughput within a sterile enclosure equipped with HEPA-filtered airflow. The Beckman Coulter software suite, comprising Biomek 5 for method development and SAMI for centralized workflow management, scheduling, and extended method design, provides a robust framework for executing comprehensive automated liquid handling workflows.

### 2. CRISPRa cell line establishment

HEK293 cells were manually co-transduced with lentiviruses expressing GFP(1-10)- VP64-NLS (a gift from Bo Huang, Addgene plasmid #70228) and NLS-dCas9-GFP11x7- NLS (a gift from Bo Huang, Addgene plasmid #70224) [3], followed by fluorescence-activated cell sorting using FACSAri III Cell Sorter (BD Biosciences) to enrich for GFP-positive cells for CRISPRa-dCas9-GFP-VP64 cell line establishment.

### 3. RT-qPCR quantification to evaluate CRISPRa efficiency

To evaluate the efficiency of the T. gonfio library for CRISPR gene activation, HEK293 cells expressing CRISPRa-dCas9-GFP-VP64 were transduced with cherry-picked qgRNAs targeting the promoters of kinase and phosphatase genes, along with non-specific qgRNAs. Based on the HPA Cell Line Gene Expression Profiles [41], we selected eight kinase and phosphatase genes that had low transcript expression in HEK293 cells compared to other lines. We included three NS qgRNAs (NS6, NS10, and NS12) to serve as negative controls for baseline gene expression. Following puromycin selection, mRNA was extracted using the RNeasy Mini Kit (Qiagen) at day 7 post-transduction. RT-qPCR was subsequently performed using the Brilliant II SYBR Master Mix (Agilent) on the LightCycler 96 instrument (Roche), according to the manufacturer’s instructions, with all steps conducted manually. PCR primer sequences were summarized in Supplementary Table 1.

### 4. Arrayed lentiviral CRISPRa library establishment

Based on previously published studies reporting the activation of nine kinases and two phosphatases implicated in promoting or restricting cancer cell proliferation, an arrayed T. gonfio mini-library consisting of these 11 kinase/phosphatase-targeting, along with 12 non-specific (NS) qgRNAs, was produced, titrated, and cryopreserved using the Biomek i7 system, following workflows described in a previous study [11]. Compared with the three NS qgRNAs (NS6, NS10, and NS12) used in the RT-qPCR assay as negative controls to evaluate CRISPRa efficiency, the library screens employed twelve non-specific qgRNAs (NS1-NS12) to better evaluate the baseline of no-effect or non-specific phenotypes resulting from CRISPRa activity. This ready-to-spot lentiviral mini-library contained 50 µL of supernatant per well per lentivector within a 96-well conical bottom plate. The last column of wells was virus-free as a negative control.

### 5. Arrayed CRISPRa library screening

The high throughput arrayed CRISPRa library screening workflow (Figure 1) developed in this study consists of three Biomek automated pipelines: 1) lentiviral library transduction, 2) cell library passaging, and 3) assay processing.

**Figure 1:**
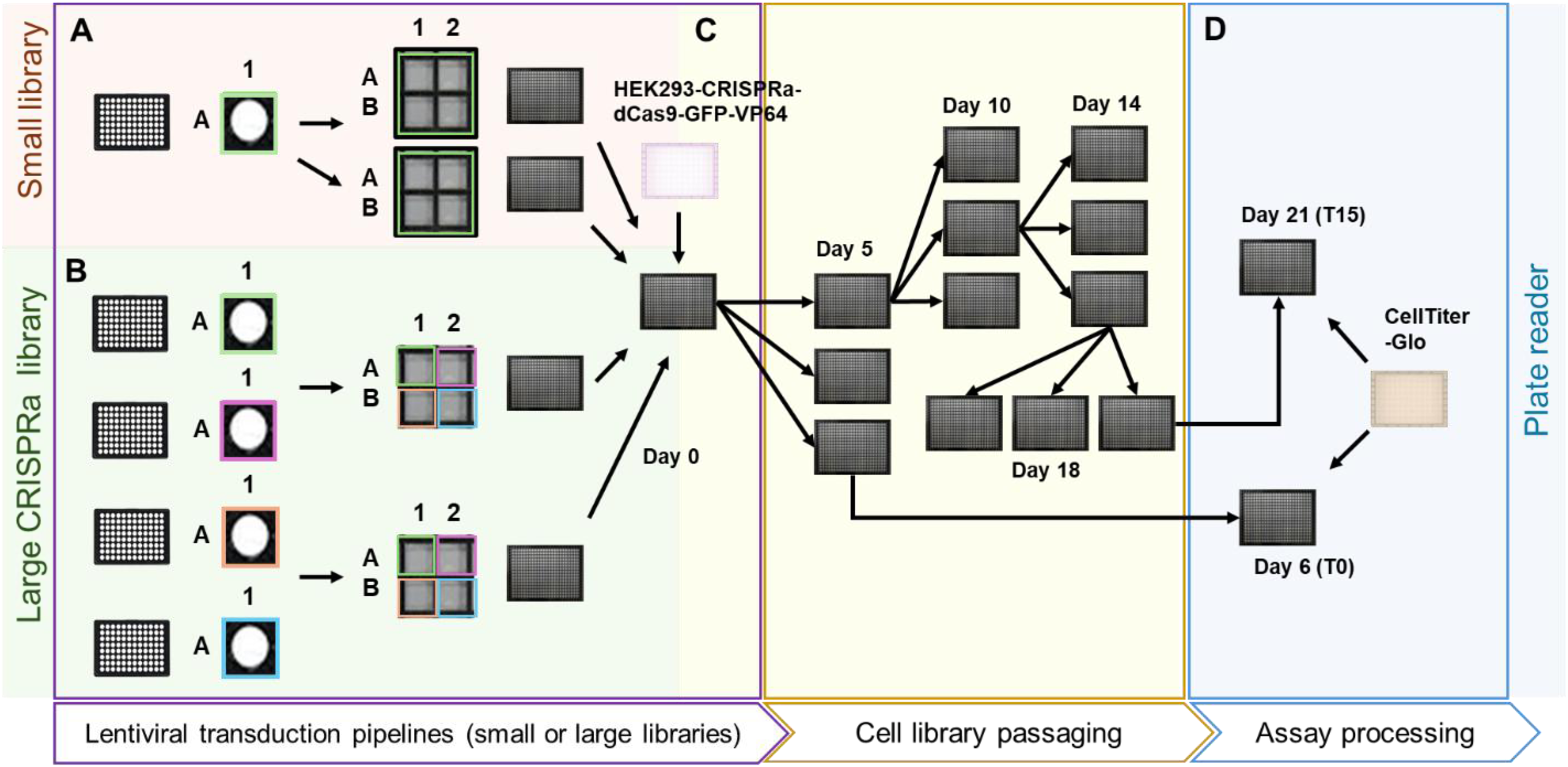
Workflow for automated arrayed CRISPRa library high-throughput screening. Schematic overview of the workflow, where one or more lentiviral libraries were freshly thawed as the starting material and processed through the entire screening workflow to yield 384-well plates with cells as the final output for plate reader analysis. The first pipeline, which involved lentivirus transduction, was described previously [11]. **A.** Briefly, for library sizes of 88 samples or fewer, lentiviruses from each well of one 96-well conical-bottom plate were transferred four times into four replicate wells of a 384-well plate. This spotting process (highlighted with an orange background) yielded two 384-well plates as biological replicates. **B.** In contrast, when the library had four 96-well plates (equivalent to 352 lentiviral samples) or more, each well of the four 96-well plates were transferred to distinct quadruplicate positions in two 384-well plates, yielding two biological replicates (highlighted with a green background). **C.** HEK293-CRISPRa-dCas9-GFP-VP64 cells were transferred to the spotted libraries from Figures 1A and 1B, and this time point was defined as day 0 post-transduction (PT) and the beginning of the second pipeline for cell library passaging (highlighted with a yellow background). At 5 days PT, the transduced cell libraries in the 384-well plate format were passaged into triplicate plates at a 1:16 dilution ratio, based on the procedures described in Materials and Methods. Of these three plates, one was analyzed using the CellTiter-Glo (CTG) assay after overnight incubation to establish the T0 time point (equivalent to 6 days PT) of the CRISPRa screen. The second plate served as a backup, and the third plate was passaged again after an additional 5 days of incubation (10 days PT). The cell library passaging pipeline was repeated every four days for two additional passages (14 and 18 days PT). **D.** Subsequently, after the last cell passage at 18 days PT and three days incubation (21 days PT), the resulting 384-well plates were analyzed via CellTiter-Glo assay using the assay processing pipeline to evaluate the CRISPRa screening results at T15 (highlighted with a blue background).

The lentiviral library transduction pipeline, which included viral library spotting and cell plate transduction, was identical to the lentivirus titration pipeline for small libraries described previously (Figure 1A) [11]. Briefly, an arrayed CRISPRa T. gonfio mini-library screen was performed in 4 biological replicate wells to evaluate promoter activation associated with growth enhancement or arrest in HEK293 cells. The lentiviral mini-library preserved in 96-well conical bottom plate was thawed. From each source well, 5 µL of virus (median titer: 10^6^ TU/mL) was transferred in quadruplicate into four replicate wells of a 384-well screening plate to maintain the original library distribution. Aliquots were used to transduce 500 HEK293-CRISPRa-dCas9-GFP-VP64 cells per well in a 384-well plate with complete DMEM (10% FBS, 1X Penicillin-Streptomycin). Transduction was performed at an average M.O.I. of 10 to achieve near 100% efficiency. At 5 days post-transduction, the screening plate underwent arrayed cell culture passaging into three 384- well plates at a 1:16. One 384-well plate with overnight incubation, serving as the normalization control at time point 0 (T0; time-point 0, equivalent to 6 days post-transduction), was immediately used to measure cell density via the CellTiter-Glo cell survival assay (Promega) and Envision 2103 luminescence plate reader (PerkinElmer) in SBP Therapeutic Discovery Core Facility. One 384-well plate was retained as a backup. The last plate was incubated at 37 °C in a Cytomat 2C incubator for an additional 5 days, followed by three additional 1:16 passaging to extend the culture period to time point 15 (T15), at which final cell growth was measured using the CellTiter-Glo assay (Figures 1C and 1D).

### 6. Arrayed cell library passaging

This pipeline (Figure 1C) was adapted and optimized from a previous study [10]. The layouts of the incubator and the liquid-handling deck are shown in Figure 2G. A parental 384-well plate containing HEK293-CRISPRa-dCas9-GFP-VP64 cells transduced with arrayed CRISPRa T. gonfio mini-library after 5 days post-transduction was transferred from the incubator to the CloneSelect Imager (CSI) using the Beckman Robotic Transport II (BRT II). The CSI measured cell confluency at high throughput in approximately 90 seconds per 384-well plate [10]. After imaging, the BRT II transported the plate through a barcode scanner for hardware tracking and data recording. A liquid-handling gripper then placed the plate on the designated deck position and removed the lid, followed by media removal using a 384-channel pipette head. After aspiration, 40 µL of 1X PBS without magnesium and calcium was gently dispensed using the same head at 1µL/sec speed, with each tip touching the side wall of the well. The entire PBS wash step was repeated twice. After the second PBS removal, 40 µL of 0.25% trypsin was added and the lid was placed back on the plate. The plate was incubated at 37 °C on a BioShake 3000-T ELM microplate shaker for 8 min and vortexing on the shaker at 600 rpm for an additional 30 sec to dissociate cells from the bottom of the well. After cell trypsinization, 40 µL of complete medium was dispensed and mixed by pipetting up and down vigorously 100 times at a speed of 300 µL/sec, with the tip positioned 0.1 mm above the well bottom at 5 different coordinates within each well (20 times each in the center, and the four corner positions). After dissociation of cell aggregates, a repetitive cell suspension stamping in 5 µL per well per plate was performed at a 1:16 ratio into triplicate daughter plates using a 384-channel pipette head. All daughter plates were pre-filled with complete media.

**Figure 2:**
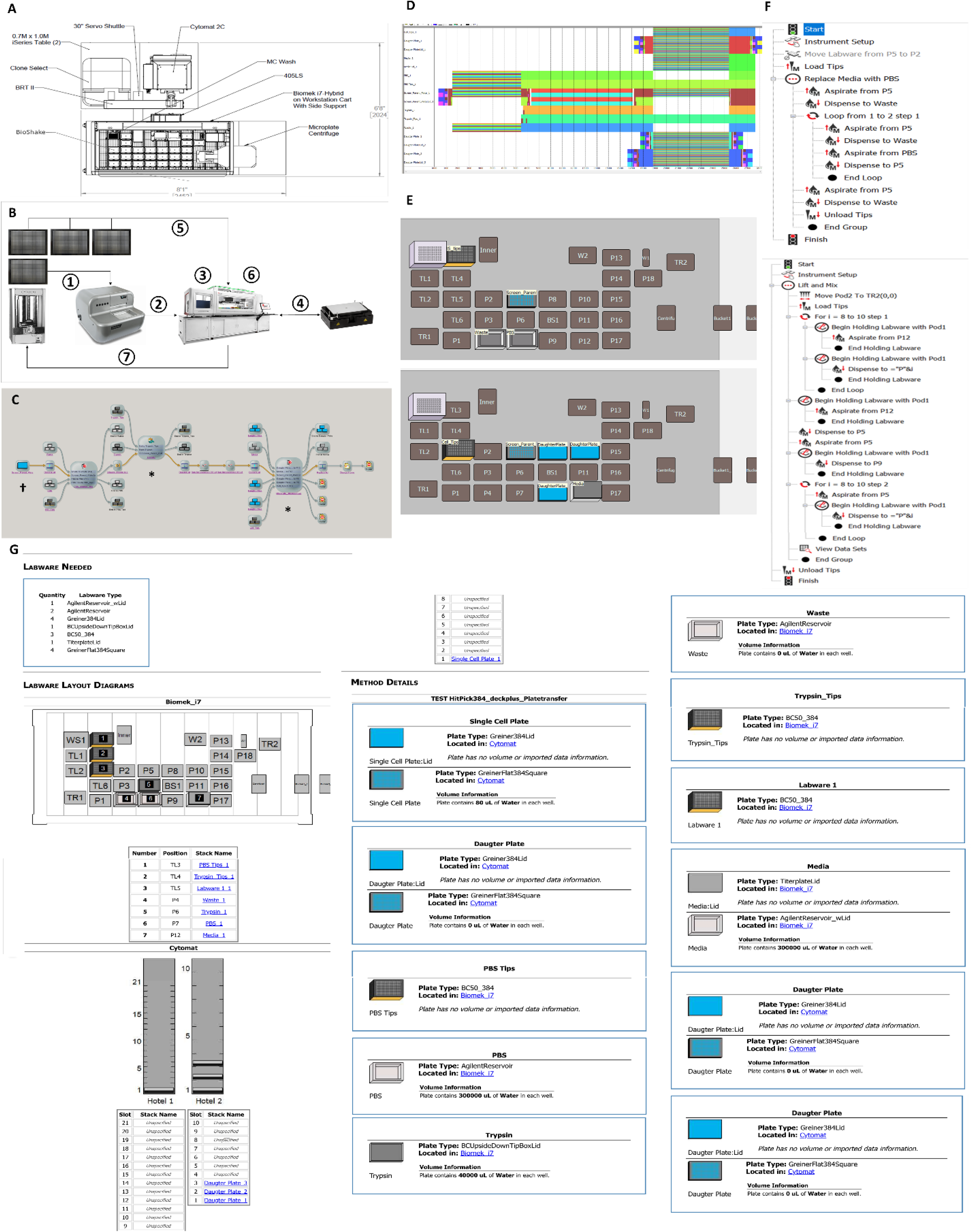
High-throughput pipeline for automated passaging of the arrayed cell library. **A.** Hardware design of the Biomek i7 Hybrid automated workstation. **B.** Schematic overview of arrayed cell library passaging. ① A parental 384-well plate containing cells transduced with the lentiviral CRISPRa min-library was transferred from the Cytomat 2C to the CSI via BRT II. ② The parental plate was transferred to the Biomek i7 deck position via the servo shuttle and track gripper. ③ The parental plate underwent liquid removal, two PBS washes, and trypsin treatment using the 384-channel pipette head. ④ Plate incubation at 37°C and vortexing during trypsinization were carried out using the BioShake. ⑤ Three new daughter plates were transferred from the Cytomat 2C incubator to the Biomek i7 deck. ⑥ Media addition and mixing during the neutralization and backfill steps, as well as cell suspension transfer, were performed using the 384-channel pipette head. ⑦ All parental and daughter plates were transferred back to the incubator. **C.** Pipeline overview under the SAMI EX interface. SAMI EX and Biomek 5 software were utilized for the method creation and pipeline design. **D.** Scheduling and time estimation for the entire pipeline by SAMI EX illustrate the timestamps of each labware item in chronological order. **E.** Deck layout of two automated Biomek 5 methods specific to PBS replacement (top panel) and cell suspension mixing (bottom panel) are shown at the position indicated by the asterisk in Figure 2C. **F.** Overview of two Biomek 5 methods for PBS replacement (top panel) and cell suspension mixing (bottom panel), indicated by the asterisk in Figure 2C. **G.** The labware setup report, marked by the dagger symbol in Figure 2C, serves as a starting reference and provides a comprehensive overview of the SAMI EX deck layout, along with details on labware and associated conditions.

#### 6.1. Calculation of suspension volumes for cell passaging

To passage cells at a defined ratio rather than based on an absolute seeding number, a user-generated CSV file, instead of an auto-exported file from the CSI, was manually uploaded to the Biomek system. This ensured a uniform transfer volume of 5 µL per well at a ratio of 1:16 across the entire plate. The equal volume transfer was calculated using the formula below for consistent seeding across wells.

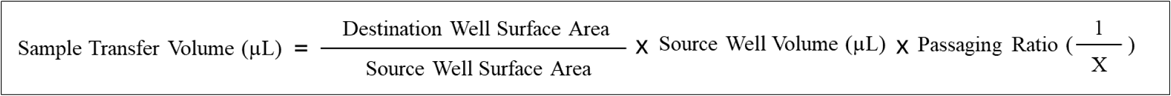

### 7. Arrayed assay processing

The layouts of the incubator and liquid-handling deck are shown in Figure 3F. Three 384- well plates, derived from four rounds of cell passaging at a 1:16 ratio (1/16 cell dilution) and 21 days of incubation post-transduction, were sequentially transferred from the Cytomat 2C to the CSI for cell density imaging and subsequently to designated positions on the Biomek deck. Plates were spun using a microplate centrifuge at 500 rpm for 1 min to pull down residual liquid. A 384-channel pipette head then aspirated 40 µL of medium at a rate of 2 µL/sec, with the tips positioned 0.1 mm from the well bottom. After partial medium removal, 30 µL of CellTiter-Glo reagent was dispensed at a rate of 150 µL/sec, with the tips positioned 0.1 mm from the well bottom. Plates were briefly centrifuged again and returned to the deck for 20 min of incubation at room temperature before bioluminescence plate reader analysis (Figure 1D).

**Figure 3:**
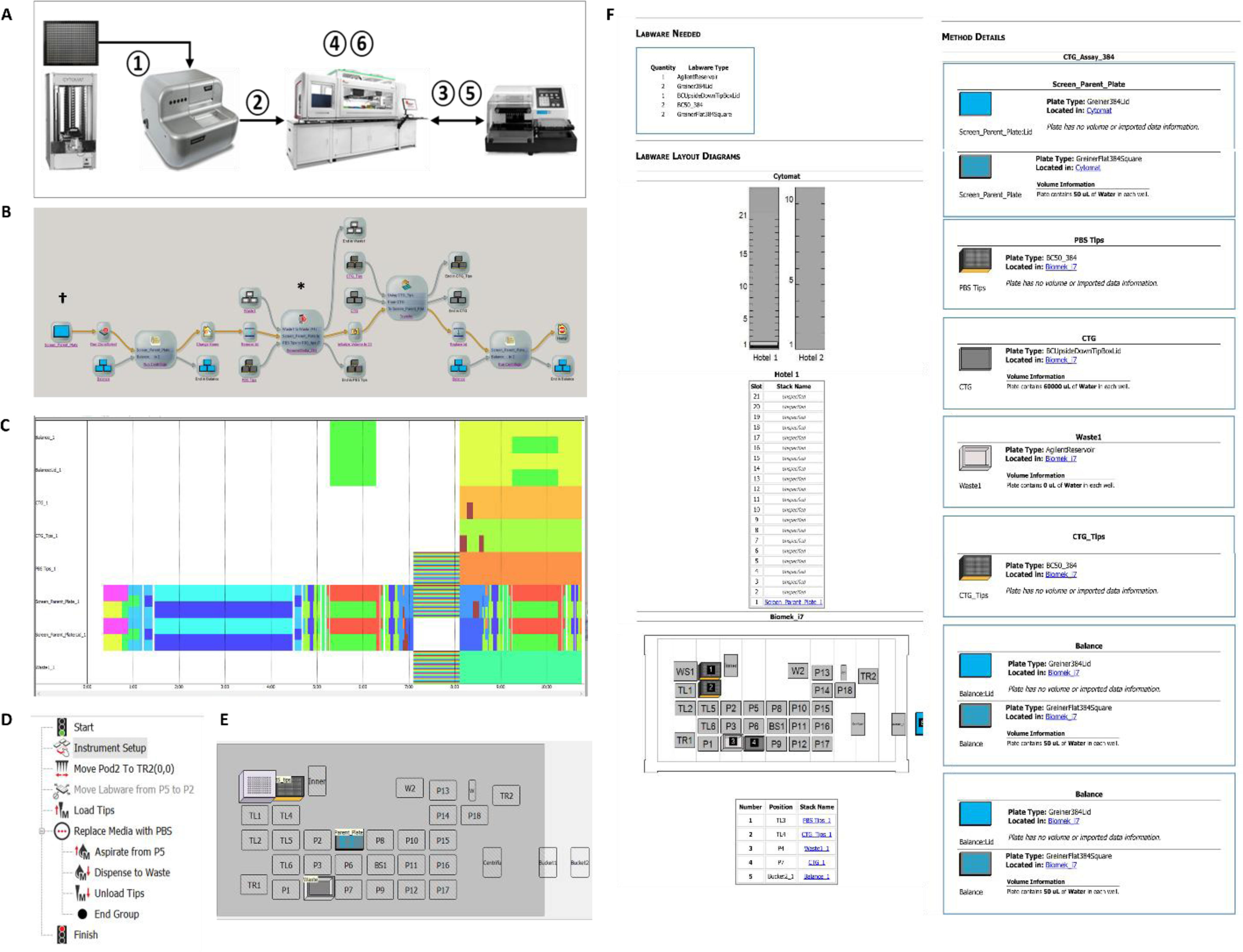
High-throughput pipeline for automated arrayed assay processing. **A.** Schematic overview of the arrayed CellTiter-Glo assay pipeline. ① Following cell passaging and incubation at 21 days post-transduction, a 384-well screening plate containing cells transduced with the lentiviral CRISPRa min-library was transferred from the Cytomat 2C to the CSI via BRT II. ② The plate was subsequently transferred to the Biomek i7 deck position using the servo shuttle and track gripper. ③ The plate underwent brief centrifugation. ④ A 384-channel pipette head partially aspirated the medium. ⑤ The plate was centrifuged again. ⑥ The plate was transferred to the Biomek i7 deck for incubation. **B.** Overview of the pipeline under the SAMI EX interface. SAMI EX and Biomek 5 software were used for method development and pipeline design. **C.** Scheduling and time estimation for the entire pipeline in SAMI EX, illustrating the chronological timestamps of each labware item. **D.** Biomek 5 method layout for partial medium removal and CellTiter-Glo lysis buffer addition. **E.** Deck layout of the Biomek 5 method for partial medium removal and CellTiter-Glo lysis buffer addition, corresponding to the position indicated by the asterisk in Figure 3B. **F.** Labware setup report, indicated by the dagger symbol in Figure 3B, providing a reference overview of the SAMI EX deck layout, including labware details and associated conditions.

### 8. Pooled CRISPRa library screening

A pooled mini-library screen consisting of 11 kinase/phosphatase and 12 non-specific qgRNAs (NS1-NS12) was conducted in 2 biological replicates to confirm promoter activation related growth phenotype in HEK293 cells. HEK293 cells expressing CRISPRa-dCas9-GFP-VP64 were transduced at an M.O.I. of 0.3 with coverage exceeding 25,000-folds above the library size. After 3 days of transduction and 2 additional days of puromycin selection, samples were determined to be at the baseline time point (T0). The cell libraries were then cultured, and 2 million cell pellets were collected for genomic DNA extraction at days 14 (T9) and 29 (T24) post-transduction. Amplicon DNA libraries were prepared from T0, T9, and T24 samples for TapeStation and next-generation sequencing analyses at the SBP Genomics Core Facility.

### 9. Arrayed deconvolution CRISPRa library screening

Four individual single-gRNAs (sgRNAs), derived from EPHA2-qgRNAs (Supplementary Table 2), were manually cloned into the sgRNA-BFP-Puro lentivector (a gift from Jonathan Weissman and Luke Gilbert). For the deconvolution screen, individual plasmid and virus encoding four sgRNAs and one qgRNA against EPHA2 were prepared separately. The experimental materials, procedures, conditions, and assay of the deconvolution screen were identical to those used in the arrayed CRISPRa library screening, except the screening format was in a 96-well rather than a 384-well plate format. At 21 days post-transduction, including four cell culture passages, cell viability was assessed using the CellTiter-Glo assay.

### 10. Cell growth curve assessment through co-culture

To validate the finding that EPHA2 promoter activation reduces cell proliferation, wild-type HEK293 were co-cultured with HEK293 cells expressing either CRISPRa-qgEPHA2-BFP or CRISPRa-qgNS-BFP cells at a 15:85 ratio, with three biological replicates per condition. Five days after transduction, the initial time point (T0) was established by mixing the newly generated CRISPRa-qgEPHA2-BFP and CRISPRa-qgNS-BFP cells with wild-type HEK293 cells. The co-cultured cells were subsequently passaged every 4-5 days at a 1:10 split ratio and the remaining cell suspension was used to monitor the percentage of BFP-positive cells using the LSRFortessa flow cytometer (BD Biosciences). In the case of a growth inhibitory phenotype, the percentage of BFP-positive cells in the CRISPRa- qgNS-BFP samples (NS10 as the negative control) was expected to remain stable at approximately 85% over time, whereas that in the CRISPRa-qgEPHA2-BFP samples was expected to be lower than 85% and to decrease progressively over time. The same experimental design was used for the co-culture assay with wild-type HEK293 cells mixed with CRISPRa-BFP-sgEPHA2-#1, #2, #3, or #4 cells.

## Result

### High-throughput automation design

The Biomek i7 Hybrid automated workstation has a dual-pod liquid handling system that supports high-throughput arrayed mammalian cell culture [10] and lentivirus production [11]. To achieve a fully automated, high-throughput arrayed CRISPRa screening workflow, the Biomek i7 platform was equipped with a left pod fitted with interchangeable 96 or 384- channel pipette heads for multi-well pipetting, and a right pod containing eight independent pipette heads for sample cherry-picking and plate reformatting. In addition to the Biomek liquid handler, multiple instruments were integrated into a unified system (Figure 2A), each contributing a distinct function to the automation workflow (Figures 2B, 3A). The Cytomat 2C tower supported cell culture incubation across various microplate formats. The CloneSelect Imager (CSI) enabled high-throughput assessment of cell confluency. The BioShake 3000-T ELM shaker offered heating and orbital mixing functions. The microcentrifuge was used to quickly clarify supernatant prior to downstream applications.

### Assessment of the CRISPRa efficiency of the T. gonfio library

The T. gonfio kinase, phosphatase, and drug-target sub-library targets 2,012 gene promoters. Prior to library screening, eight cherry-picked genes from this sub-library were used to evaluate CRISPRa efficiency in the HEK293-CRISPRa-dCas9-GFP-VP64 cell model. A total of eight genes, including seven kinases (CSNK1A1L, EPHA2, GSK3B, MARK3, POMK, RIOK2, and ROCK1) and one phosphatase (PTPRO) were selected based on their low mRNA expression in HEK293 cells relative to other cell lines, according to HPA Cell Line Gene Expression Profiles [41]. Three non-specific (NS) qgRNAs, including NS6, NS10, and NS12, served as negative controls. Seven days after transduction, six genes (CSNK1A1L, EPHA2, GSK3B, MARK3, PTPRO and RIOK2) were found to be successfully upregulated at the mRNA level, demonstrating robust CRISPRa efficiency (Figure 4A). Among these, CSNK1A1L, EPHA2, MARK3 and PTPRO were confirmed to have low endogenous basal transcript levels, as indicated by relatively higher baseline Ct values (27-30) in RT-qPCR assays (data not shown). Of the remaining four genes with higher basal mRNA levels (Ct values 19-22; data not shown), transcription of POMK and ROCK1 was not detectably activated.

**Figure 4:**
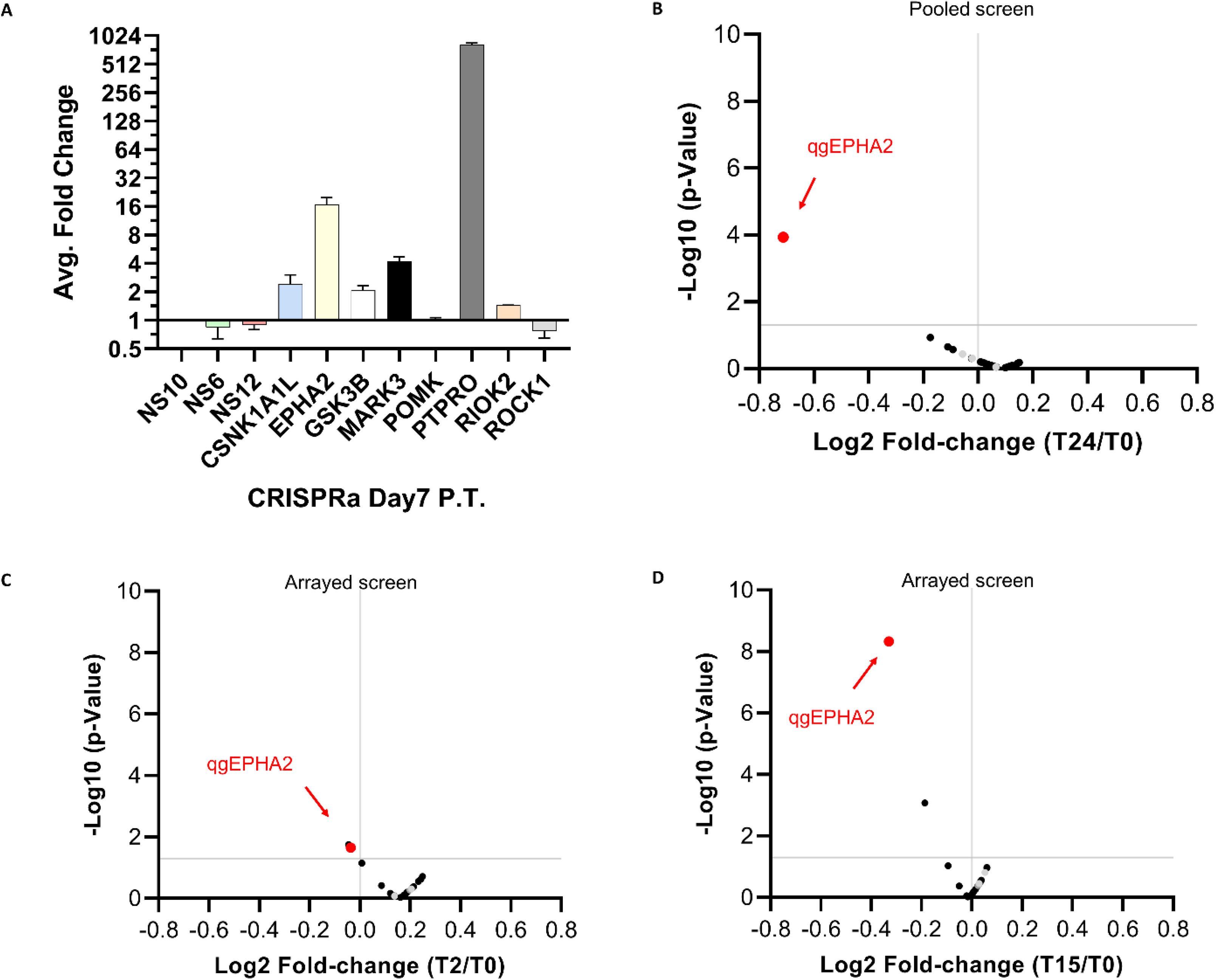
Evaluation of the T. gonfio library for CRISPR gene activation in pooled and arrayed screens. **A.** RT-qPCR quantification of mRNA levels for eight genes at day 7 post-transduction (PT) of CRISPRa-qgRNAs in HEK293 cells. NS: non-specific qgRNA. **B.** Volcano plot of the pooled mini-library screen showing log2 fold change in cell viability (x-axis) versus -log10 adjusted *P*-value (y-axis) for all qgRNAs, including 11 kinase- or phosphatase-targeting and 12 non-specific qgRNAs. The horizontal gray line indicates the significance threshold (P = 0.05). Cell viability was assessed based on qgRNA reads obtained from next-generation sequencing as the readout for pooled screens. The EPHA2-targeting qgRNA that significantly downregulated cell growth is highlighted in red. Non-specific qgRNA-#6, #10, and #12 samples are highlighted in gray. Each qgRNA was tested in two biological replicates. Day 5 PT was designated as time point 0 (T0), and T24 corresponded to day 29 PT. **C.** Volcano plot of the arrayed mini-library screen at day 7 PT (T2), with eight biological replicates per qgRNA (T0 = day 5 PT). **D.** Volcano plot of the same arrayed mini-library screen at day 21 PT (T15), with eight biological replicates per qgRNA (T0 = day 6 PT). Cell viability in panels C and D was assessed using the CellTiter-Glo assay as the readout for arrayed screens.

### Arrayed CRISPRa library screening workflow

The high-throughput arrayed CRISPRa library screening workflow can be scaled to larger libraries (Figure 1B). To validate the workflow and assess library screening efficiency, HEK293-CRISPRa-dCas9-GFP-VP64 cells were used as a highly proliferative, non-cancerous cell model. The CRISPRa T. gonfio mini-library containing 11 kinase- or phosphatase-targeting qgRNAs was used to evaluate the effects of candidate gene promoter activation on cell growth. Twelve non-specific qgRNAs served as non-targeting controls [5]. The first pipeline of the library screening workflow, arrayed lentiviral library transduction, used the same layout and methods to prepare the lentiviral T. gonfio mini-library from three 96-well virus-producing plates [11] and to transfer viruses from an aliquoted 96-well storage plate to two 384-well screening plates (Figure 1A), generating a quadruplicate virus-transduced cell library in approximately one hour (Table 1) [11]. Thus, a total of eight biological replicate wells were included per qgRNA to increase the statistical power of the library screen.

**Table 1.**
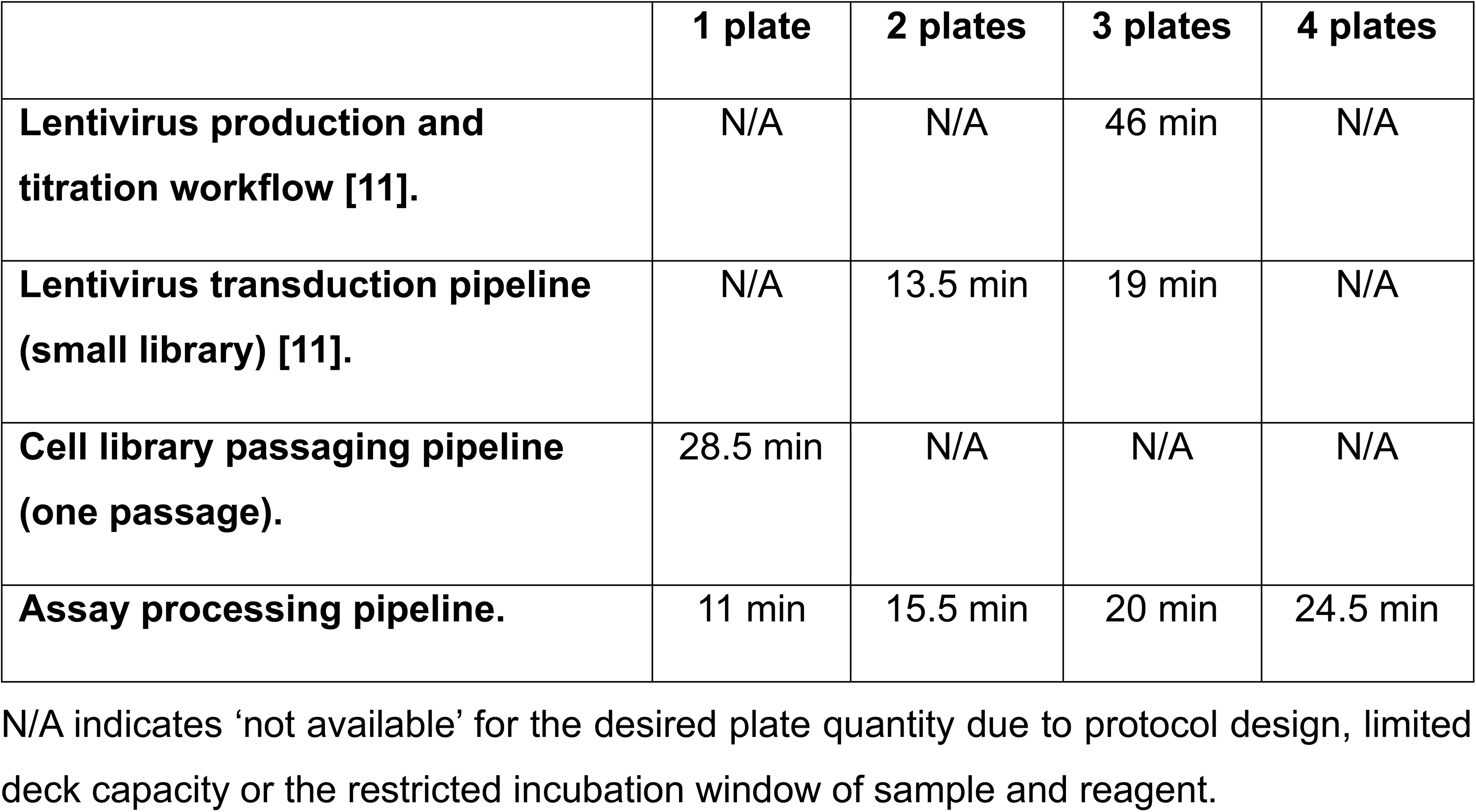
Time required for each automation pipeline.

|  | **1 plate** | **2 plates** | **3 plates** | **4 plates** |
| --- | --- | --- | --- | --- |
| **Lentivirus production and titration workflow [11].** | N/A | N/A | 46 min | N/A |
| **Lentivirus transduction pipeline (small library) [11].** | N/A | 13.5 min | 19 min | N/A |
| **Cell library passaging pipeline (one passage).** | 28.5 min | N/A | N/A | N/A |
| **Assay processing pipeline.** | 11 min | 15.5 min | 20 min | 24.5 min |

N/A indicates ‘not available’ for the desired plate quantity due to protocol design, limited deck capacity or the restricted incubation window of sample and reagent.

### Arrayed cell library passaging pipeline

Unlike the arrayed mammalian cell culture workflow, which passages cells from a single 96-well plate, and the arrayed monoclonality screening workflow, which passages cells from a 384-well plate to a 96-well plate [10], our arrayed cell library passaging pipeline was specifically designed for cell passaging within a 384-well plate format (Figures 1C, 2B) and relies on surface area ratio rather than cell seeding number (Materials and Methods 6.1). Both Biomek 5 and SAMI EX software programs were used to develop methods integrated into a unified pipeline controlled by the SAMI interface (Figure 2C). Beginning with the labware setup report, which summarizes the initial deck layout and labware details (Figure 2G), SAMI EX can function as the master controller for scheduling, simulation, multitasking, and time estimation of the entire pipeline (Figure 2D). Using distinct deck layouts (Figure 2E), two Biomek 5 methods executed key pipetting steps, including liquid removal, PBS washes, trypsinization, neutralization, and cell disaggregation (Figure 2F), as indicated by two asterisk symbols in Figure 2C. The runtime passage of one 384-well plate into three 384-well plates was 28.5 minutes (Figure 2D, Table 1).

### Arrayed assay processing pipeline

Four rounds of cell library passaging using the arrayed cell passaging pipeline (Figure 1C) extended the duration of the arrayed CRISPRa screening to time point 15 (T15, 21 days post-transduction). The 384-well screening plates were processed through this pipeline for assay readout and analysis at 6 days post-transduction (corresponding to T0) and at T15 (Figure 1D). SAMI EX was also used for pipeline design (Figure 3A) and method development (Figure 3B), except for a liquid-handling step in which a 384-channel pipette head was used to aspirate medium and dispense lysis buffer for the CellTiter-Glo assay. The method (Figure 3D) and corresponding deck configuration (Figure 3E) for this pipetting step were implemented using Biomek 5 software, as indicated by the asterisk in Figure 3B. The labware setup report, denoted by the dagger symbol in Figure 3B, summarizes the initial deck layout, labware, and associated conditions (Figure 3F). SAMI EX scheduled and processed one to four 384-well plates in 11 to 25 min (Figure 3C, Table 1).

### Activation of the EPHA2 promoter inhibits cell growth

Both arrayed and pooled CRISPRa screens are suitable for assessing how candidate gene promoter activation influences cell growth. The CellTiter-Glo assay was used to quantify viable cells as the readout of the screens. Nine kinase genes (including AKT1, CDK1, CDK2, CDKL3, EPHA2, JAK2, MAP2K1, MAPK1, and PLK1) and the PPP2R1A phosphatase have been reported to promote cancer cell proliferation upon activation [42, 43, 44, 45, 46, 47, 48, 49, 50]. However, ligand-induced EPHA2 tyrosine kinase activity can inhibit the growth of normal human mammary epithelial cells [56, 59, 61]. In contrast, PPP2CA phosphatase activation has been reported to reduce cancer cell proliferation [51, 52]. In the initial arrayed CRISPRa screen with the T. gonfio mini-library, EPHA2 was identified as a hit after 21 days of CRISPR activation (21 days post-transduction). CRISPRa-mediated EPHA2 upregulation led to a statistically significant 20% reduction in cell growth from day 6 post-transduction (T0) to day 21 post-transduction (T15; Figure 4D). A pooled CRISPRa mini-library screen also confirmed the dropout of EPHA2 qgRNAs as a hit, showing a growth reduction phenotype of around 40% compared with qgRNAs targeting other genes that might affect cell growth (including AKT1, CDK1, CDK2, CDKL3, JAK2, MAP2K1, MAPK1, PLK1, PPP2R1A, and PPP2CA) and non-specific controls (NS1-NS12) at 29 days post-activation (Figure 4B).

To further validate the phenotype using a co-culture assay, wild-type HEK293 cells were mixed with either CRISPRa-qgEPHA2-BFP or CRISPRa-qgNS10-BFP cells at a 15% WT cells to 85% CRISPRa-BFP cells ratio. The percentage of blue fluorescent protein (BFP) positive cells was monitored every 4-5 days for nearly two months post-transduction. As shown in Figure 5A, the percentage of CRISPRa-qgEPHA2 cells gradually declined, accompanied by a corresponding increase in the percentage of co-cultured wild-type cells. After normalizing the percentage of control CRISPRa-qgNS10-BFP from 85% to 100%, cells with EPHA2 upregulation showed more than a 30% reduction in abundance compared with control cells at T52 (57 days post-transduction). Consistent with the arrayed screen results, cells transduced with CRISPRa-qgEPHA2 exhibited approximately a 20% reduction in growth at 21 days post-transduction (Figures 3D and 5A).

**Figure 5:**
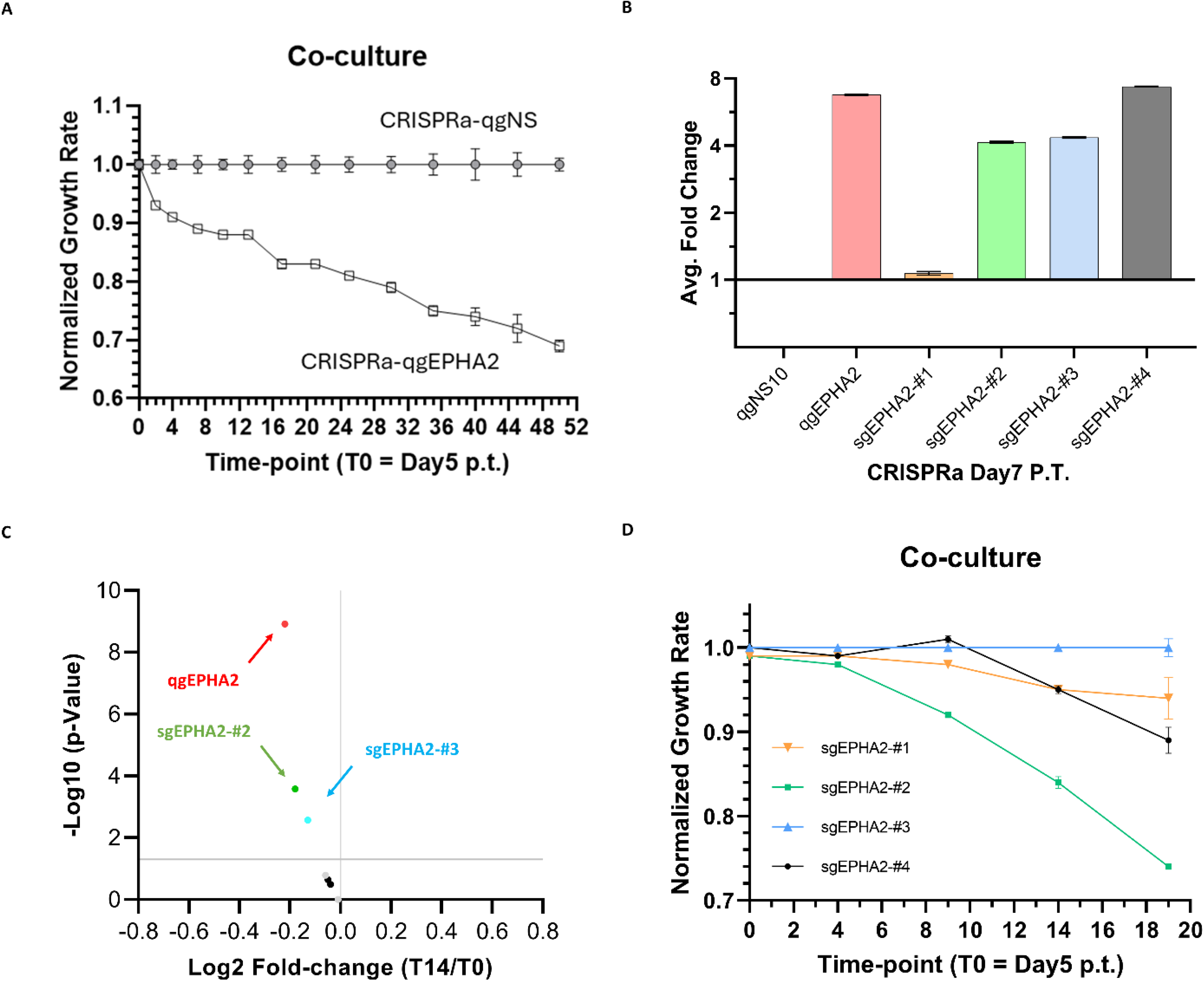
Deconvolution of the arrayed CRISPRa screening qgRNAs targeting EPHA2. **A.** Confirmation of EPHA2 promoter activation-mediated growth inhibition by comparing HEK293-CRISPRa-qgNS-BFP/HEK293-wt and HEK293-CRISPRa-qgEPHA2-BFP/HEK293-wt co-cultures (wt, wild type; n = 3 per condition). Flow cytometry sampling was performed every 4–5 days during passaging. Initial co-culture ratio: 85:15. BFP% from HEK293-CRISPRa-qgEPHA2-BFP/HEK293-wt was normalized to HEK293-CRISPRa-qgNS-BFP/HEK293-wt to indicate relatively reduced CRISPRa-qgEPHA2 cell growth. Time point 0 (T0) corresponds to 5 days post-transduction of CRISPRa-qgRNAs. **B.** RT-qPCR quantification of EPHA2 mRNA levels at day 7 post-transduction of CRISPRa-qgEPHA2 and individual CRISPRa-sgEPHA2-#1 to -#4 in HEK293 cells (qg, quadruple gRNA; sg, single gRNA). qgNS10: non-specific qgRNA-#10. **C.** Volcano plot of the arrayed deconvolution screen at day 21 PT (T15), with eight replicates per qgRNA and sgRNA against EPHA2 (T0 = day 6 PT). Cell viability was assessed using the CellTiter-Glo assay. Non-specific qgRNA-#6 and -#10 are highlighted in gray. **D.** Confirmation of EPHA2 promoter activation–mediated growth inhibition by comparing HEK293-CRISPRa-BFP-sgEPHA2 (#1, #2. #3, and #4) cells, cocultured with HEK293-wt cells and analyzed by flow cytometry every 4-5 days (n = 3 per condition).

### CRISPRa deconvolution screen

After confirming that CRISPRa-qgEPHA2 contributed to the growth phenotype, a deconvolution screen was designed to determine the contribution of the four individual single gRNAs (sgRNA), including sgEPHA2-#1, #2, #3, and #4, to the phenotype. Except for the sgEPHA2-#1, three of the four individual gRNAs targeting EPHA2 upregulated EPHA2 mRNA levels. sgEPHA2-#4 induced an 8-fold EPHA2 upregulation, similar to the EPHA2 qgRNA, while sgEPHA2-#2 and #3 induced a 4-fold upregulation (Figure 5B). Both qgEPHA2 and sgEPHA2-#2 exhibited a similar and statistically significant reduction in cell growth from day 6 post-transduction (T0) to day 21 (Figure 5C). Growth inhibition by sgEPHA2-#3 was also significant but less pronounced, while sgEPHA2-#1 and sgEPHA2-#4 showed no significant effect (Figure 5C). The co-culture assay further confirmed that sgEPHA2-#2 was a valid hit reproducing the growth inhibition phenotype of qgEPHA2 (Figure 5D). No significant effect was observed for sgEPHA2-#3 (Figure 5D).

## Discussion

Among academic resources, the T. gonfio library represents the only genome-wide arrayed CRISPR activation library in which each well contains qgRNAs targeting each gene. This design, featuring four tandem Cas9 gRNAs per lentivector, results in a smaller library size [5]. Compared with single- or dual-gRNA strategies, the distribution of four gRNAs across different promoter regions synergistically enhances both the likelihood and the efficiency of promoter activation and thus target gene expression. The present study first validated the efficiency of qgRNAs from the T. gonfio library, with an overall success rate of 75% for increased target gene expression, as determined by RT-qPCR. Notably, a 100% activation rate was observed for genes with low endogenous expression and a 50% activation rate for genes with higher basal expression.

For most immortalized adherent cell models, the effective arrayed screening window is often limited to less than one week, as cultures seeded at low densities of 250-500 cells per well in a 384 well plate rapidly reach over-confluency. This study confirms that CRISPR promoter activation was achieved by 7 days post-transduction (Figure 4A). However, no clear EPHA2-associated growth phenotype was observed within this restricted timeframe (Figure 4C). In contrast, extending the assay time by passaging the library of CRISPRa cells allowed the screen to continue until day 21 post-transduction (T15), where the EPHA2 hit was clearly identifiable (Figure 4D). Our results show that passaging allows for longer screening windows, improving the chance of detecting phenotypes that require more time to develop. To meet the need for extended arrayed CRISPR library screening, this study developed a high throughput automated workflow on the Biomek i7 Hybrid platform, integrated with multiple instruments. Automated CRISPRa screening with HEK293 cells using the selected T. gonfio mini-library robustly and reproducibly identified EPHA2 as a hit in both arrayed and pooled formats. Activation of the EPHA2 promoter leading to reduced cell growth was further confirmed through co-culture experiments. Conversely, promoter activation of AKT1, CDK1, CDK2, CDKL3, JAK2, MAP2K1, MAPK1, PLK1, PPP2R1A, and PPP2CA had no effect on cell growth.

EPHA2, a receptor tyrosine kinase of the Eph family, is expressed at low levels in most normal adult tissues but is highly expressed in a wide range of cancers [53, 54, 58, 61].

EPHA2 can signal through both ephrin-A ligand–induced, kinase-dependent canonical pathways and noncanonical pathways that depend on EPHA2 phosphorylation on S897 by AKT, RSK and PKA [61]. While EPHA2 non-canonical signaling has been linked to pro-oncogenic effects, EPHA2 canonical signaling can inhibit cell growth, for example through inhibition of the AKT and ERK pathways [61, 62, 63, 64, 65]. EPHA2 interaction with the five ephrinA ligands, which are anchored on the cell surface through a glycosylphosphatidylinositol group, typically occurs across cell-cell junctions. Thus, the EPHA2 receptor upregulated in HEK293 cells could be activated by endogenously expressed ephrinA ligands leading to canonical signaling and inhibition of cell growth. Alternatively, or in addition, overexpression of EPHA2 in HEK293 cells can cause ligand-independent dimerization leading to cross-phosphorylation of EPHA2 receptors and activation of canonical signaling [66, 67]. In normal epithelial cells, EPHA2 canonical signaling has been shown to inhibit growth and promote differentiation [56, 59, 61, 62, 65, 68]. In normal melanocytes, EPHA2 overexpression has also been shown to trigger apoptosis through activation of a caspase-8 dependent pathway [57, 60], which could be another mechanism inhibiting cell growth. Interestingly, the EPHA2 promoter can be activated by DNA damage, for example induced by UV exposure, underscoring a role in stress-responsive signaling networks [53, 60]. In addition, the EPHA2 promoter can be activated by the ERK MAP kinase pathway in cancer cells, concomitant with decreased ephrinA expression, leading to decreased canonical signaling and pro-oncogenic effects [64, 69]. Notably, our study employed CRISPRa to engineer the immortalized HEK293 cell line in order to investigate whether promoter activation of individual kinase or phosphatase genes can influence cell proliferation, even if the phenotype is modest and thus only becomes apparent over prolonged times. Given the low endogenous EPHA2 expression, the growth reduction phenotype we observed indicates that EPHA2 promoter activation can exert antiproliferative effects in the HEK293 epithelial cell model.

The goal of the present study was to establish an automated workflow on the Biomek i7 platform for high-throughput screening of an arrayed CRISPR library (Figure 1). During this process, a growth inhibition phenotype was observed upon EPHA2 promoter activation, warranting further mechanistic investigation into the upstream and downstream regulators of this phenotype. For instance, although qgEPHA2 and sgEPHA2-#2, #3, and #4 increased EPHA2 mRNA expression (Figure 5B), only sgEPHA2-#2 and qgEPHA2 cells exhibited a significant growth inhibition phenotype in co-culture assays (Figures 5A and 5D). Growth curve analyses using cDNA-based overexpression of EPHA2 could ultimately confirm whether the phenotype is due to EPHA2 protein overexpression. Furthermore, increased expression of an ephrinA ligand, or expression of EPHA2 mutants deficient in ligand binding or kinase activity, could determine whether the growth inhibitory phenotype depends on EPHA2 canonical signaling. Alternatively, expression of the EPHA2 S897A mutant deficient in S897 phosphorylation could determine whether changes in non-canonical signaling are involved. According to the published information on the CRISPRa library [5], sgEPHA2- #2, located 154 base pairs upstream of the EPHA2 transcription start site, exhibited the highest predicted score among the four sgRNAs and had <10 GuideScan-predicted off-targets. Future studies evaluating potential off-target activity or accessory gene expression induced by sgEPHA2-#2–mediated promoter activation near the transcription start site could help elucidate the mechanism underlying the observed cell growth inhibition. In parallel, whole-genome transcriptomic profiling and proteomics / phosphoproteomics analyses of HEK293 cells transduced with CRISPRa qgRNAs and sgEPHA2-#2 targeting EPHA2 may reveal transcriptional programs and signaling pathways that are differentially regulated in response to EPHA2 promoter activation leading to the growth inhibitory phenotype.

## Acknowledgement

CH, RM, and CY were supported by the Sanford Burnham Prebys (SBP) NCI Cancer Center Support Grant P30 CA030199. Research reported in this publication was supported by the SBP Functional Genomics Core through NIH Shared Instrumentation Grant S10 OD036254, and by the SBP Flow Cytometry Core through Grant S10 OD032325. Additional support was provided through NIH grant R01 AG071861 to PDA and R01 CA262794 to EBP. We would like to acknowledge the SBP Genomics Core, and the Beckman Coulter team, including Brandon K. Corbin, Kenzo Maetani, Michael D. Moran, Eugene B. Tupas, and Liz Chladny, for their support with hardware. We would also like to acknowledge the Beckman software team, including Christopher J. Knuff, Joshua M. Yoder, Amy V. Gibson, Marc A. Post, Zarina K. Waqar, and Daniel Lynch, for their software support.

## Author contributions

CH, AB, EBP, AJD, MJ, and PDA designed the studies. CY, YW, and JAY developed the software and automation workflows. CY, RM, and CH performed the experiments and analyzed data. CH, AB, EBP, and CY wrote the manuscript.

## Conflict of Interest

Anna Beketova is an employee of Beckman Coulter Life Sciences. Other authors declare no competing financial interests.

## Reference

1. Joung J, Konermann S, Gootenberg JS, Abudayyeh OO, Platt RJ, Brigham MD, Sanjana NE, Zhang F. Genome-scale CRISPR-Cas9 knockout and transcriptional activation screening. Nat Protoc. 2017 Apr;12(4):828–863. doi: 10.1038/nprot.2017.016. Epub 2017 Mar 23. Erratum in: Nat Protoc. 2019 Jul;14(7):2259. doi: 10.1038/s41596-018-0063-0. PMID: 28333914; PMCID: PMC5526071.

2. Horlbeck MA, Gilbert LA, Villalta JE, Adamson B, Pak RA, Chen Y, Fields AP, Park CY, Corn JE, Kampmann M, Weissman JS. Compact and highly active next-generation libraries for CRISPR-mediated gene repression and activation. Elife. 2016 Sep 23;5:e19760. doi: 10.7554/eLife.19760. PMID: 27661255; PMCID: PMC5094855.

3. Kamiyama D, Sekine S, Barsi-Rhyne B, Hu J, Chen B, Gilbert LA, Ishikawa H, Leonetti MD, Marshall WF, Weissman JS, Huang B. Versatile protein tagging in cells with split fluorescent protein. Nat Commun. 2016 Mar 18;7:11046. doi: 10.1038/ncomms11046. PMID: 26988139; PMCID: PMC4802074.

4. Replogle JM, Bonnar JL, Pogson AN, Liem CR, Maier NK, Ding Y, Russell BJ, Wang X, Leng K, Guna A, Norman TM, Pak RA, Ramos DM, Ward ME, Gilbert LA, Kampmann M, Weissman JS, Jost M. Maximizing CRISPRi efficacy and accessibility with dual-sgRNA libraries and optimal effectors. Elife. 2022 Dec 28;11:e81856. doi: 10.7554/eLife.81856. PMID: 36576240; PMCID: PMC9829409.

5. Yin JA, Frick L, Scheidmann MC, Liu T, Trevisan C, Dhingra A, Spinelli A, Wu Y, Yao L, Vena DL, Knapp B, Guo J, De Cecco E, Ging K, Armani A, Oakeley EJ, Nigsch F, Jenzer J, Haegele J, Pikusa M, Täger J, Rodriguez-Nieto S, Bouris V, Ribeiro R, Baroni F, Bedi MS, Berry S, Losa M, Hornemann S, Kampmann M, Pelkmans L, Hoepfner D, Heutink P, Aguzzi A. Arrayed CRISPR libraries for the genome-wide activation, deletion and silencing of human protein-coding genes. Nat Biomed Eng. 2025 Jan;9(1):127–148. doi: 10.1038/s41551-024-01278-4. Epub 2024 Dec 4. PMID: 39633028; PMCID: PMC11754104.

6. Kim HS, Lee K, Kim SJ, Cho S, Shin HJ, Kim C, Kim JS. Arrayed CRISPR screen with image-based assay reliably uncovers host genes required for coxsackievirus infection. Genome Res. 2018;28(6):859–68. Epub 2018/05/02. doi: 10.1101/gr.230250.117. PubMed PMID: 29712754; PMCID: PMC5991512.

7. So RWL, Chung SW, Lau HHC, Watts JJ, Gaudette E, Al-Azzawi ZAM, Bishay J, Lin LT, Joung J, Wang X, Schmitt-Ulms G. Application of CRISPR genetic screens to investigate neurological diseases. Mol Neurodegener. 2019;14(1):41. Epub 2019/11/16. doi: 10.1186/s13024-019-0343-3. PubMed PMID: 31727120; PMCID: PMC6857349.

8. Heynen-Genel S, Pache L, Chanda SK, Rosen J. Functional genomic and high-content screening for target discovery and deconvolution. Expert Opin Drug Discov. 2012;7(10):955–68. Epub 2012/08/07. doi: 10.1517/17460441.2012.711311. PubMed PMID: 22860749; PMCID: PMC3954969.

9. Fennell M, Xiang Q, Hwang A, Chen C, Huang CH, Chen CC, Pelossof R, Garippa RJ. Impact of RNA-guided technologies for target identification and deconvolution. J Biomol Screen. 2014;19(10):1327–37. Epub 2014/08/29. doi: 10.1177/1087057114548414. PubMed PMID: 25163683.

10. Yang CC, Deshpande A, Jackson M, Adams PD, Lynch D, Gibson AV, Waqar ZK, Beketova A, Yin JA, Huang CT. Automation of high-throughput arrayed mammalian cell line cultivation. bioRxiv 2025.10.03.676043. doi: 10.1101/2025.10.03.676043.

11. Yang CC, Deshpande A, Jackson M, Adams PD, Altman Y, Yin JA, Wu Y, Post MA, Beketova A, Huang CT. Automation of high-throughput arrayed lentivirus production and titration. bioRxiv 2025.10.04.680488. doi: 10.1101/2025.10.04.680488.

12. Lehmann R, Severitt JC, Roddelkopf T, Junginger S, Thurow K. Biomek Cell Workstation: A Variable System for Automated Cell Cultivation. SLAS Technology. 2016;21(3):439–50. doi: 10.1177/2211068215599786.

13. Kowalski MP, Yoder A, Liu L, Pajak L. Controlling embryonic stem cell growth and differentiation by automation: enhanced and more reliable differentiation for drug discovery. J Biomol Screen. 2012;17(9):1171–9. Epub 2012/08/17. doi: 10.1177/1087057112452783. PubMed PMID: 22895460.

14. Namatame I, Ishii K, Shin T, Shimojo D, Yamagishi Y, Asano H, Kishimoto Y, Fuse H, Nishi Y, Sakurai H, Nakahata T, Sasaki-Iwaoka H. Screening Station, a novel laboratory automation system for physiologically relevant cell-based assays. SLAS Technol. 2023. Epub 2023/05/01. doi: 10.1016/j.slast.2023.04.002. PubMed PMID: 37121549.

15. Yu C, Caothien R, Jackson M, Nakao B, Pham A, Tam L, Roose-Girma M. Advanced Technologies and Automation in mES Cell Workflow. Methods Mol Biol. 2023;2631:183–206. Epub 2023/03/31. doi: 10.1007/978-1-0716-2990-1_7. PubMed PMID: 36995668.

16. Lehmann R, Gallert C, Roddelkopf T, Junginger S, Thurow K. Biomek Cell Workstation: A Flexible System for Automated 3D Cell Cultivation. J Lab Autom. 2016;21(4):568–78. Epub 2015/07/24. doi: 10.1177/2211068215594580. PubMed PMID: 26203054.

17. Ohta A, Kawai S, Pretemer Y, Nishio M, Nagata S, Fuse H, Yamagishi Y, Toguchida J. Automated cell culture system for the production of cell aggregates with growth plate- like structure from induced pluripotent stem cells. SLAS Technol. 2023 Dec;28(6):433–441. doi: 10.1016/j.slast.2023.08.002. Epub 2023 Aug 9. PMID: 37562511.

18. Raulo R, Heuson E, Siah A, Phalip V, Froidevaux R. Innovative microscale workflow from fungi cultures to Cell Wall-Degrading Enzyme screening. Microb Biotechnol. 2019;12(6):1286–92. Epub 2019/04/22. doi: 10.1111/1751-7915.13405. PubMed PMID: 31006173; PMCID: PMC6801129.

19. Abulwerdi F, Fatehi F, Manfield IW, Le Grice SFJ, Schneekloth JS, Twarock R, Stockley PG, Patel N. Dataset of high-throughput ligand screening against the RNA Packaging Signals regulating Hepatitis B Virus nucleocapsid formation. Data Brief. 2022;42:108206. Epub 2022/05/07. doi: 10.1016/j.dib.2022.108206. PubMed PMID: 35516001; PMCID: PMC9065705.

20. Griffiths RL, Berg JD. Automation of the Whole-Blood Thiopurine S-Methyltransferase (TPMT) Phenotyping Assay Using the Biomek NX(P) and Biomek i5 Liquid-Handling Workstations. SLAS Technol. 2021;26(5):488–97. Epub 2021/04/30. doi: 10.1177/24726303211008856. PubMed PMID: 33913342.

21. Terrell KA, Sempowski GD, Macintyre AN. Development and validation of an automated assay for antidrug-antibodies in rat serum. SLAS Technol. 2023. Epub 2023/04/30. doi: 10.1016/j.slast.2023.04.001. PubMed PMID: 37120133.

22. Raghavendra Achar VG, Barde SP, Mallya MV, Awasthy D, Narayan C. Optimization of Compound Plate Preparation to Address Precipitation Issue in Mammalian A549 Cytotoxicity Assay. J Lab Autom. 2016;21(3):423–31. Epub 2015/07/18. doi: 10.1177/2211068215594768. PubMed PMID: 26185254.

23. Gunderson S. The physician’s role in organ donation and the collaborative approach to potential donor families. S D J Med. 1999;52(1):9–11. Epub 1999/02/02. PubMed PMID: 9926726.

24. Bach A, Fleischer H, Wijayawardena B, Thurow K. Optimization of Automated Sample Preparation for Vitamin D Determination on a Biomek i7 Workstation. SLAS Technol. 2021;26(6):615–29. Epub 2021/07/21. doi: 10.1177/24726303211030291. PubMed PMID: 34282678.

25. Arif-Lusson R, Agabriel C, Carsin A, Cabon I, Senechal H, Poncet P, Vitte J, Busnel JM. Streamlining basophil activation testing to enable assay miniaturization and automation of sample preparation. J Immunol Methods. 2020;481–482:112793. Epub 2020/05/11. doi: 10.1016/j.jim.2020.112793. PubMed PMID: 32387696.

26. Pumford AD, Arul AB, Ford KI, Robinson RAS. Automation of On-Resin Enrichment of S-Nitrosylated Proteins for Oxidized Cysteine-Selective cPILOT. Vanderbilt Undergrad Res J. 2021;11:43–51. Epub 2022/05/27. doi: 10.15695/vurj.v11i1.5096. PubMed PMID: 35615079; PMCID: PMC9129232.

27. Sobreira TJP, Avramova L, Szilagyi B, Logsdon DL, Loren BP, Jaman Z, Hilger RT, Hosler RS, Ferreira CR, Koswara A, Thompson DH, Cooks RG, Nagy ZK. High- throughput screening of organic reactions in microdroplets using desorption electrospray ionization mass spectrometry (DESI-MS): hardware and software implementation. Anal Methods. 2020;12(28):3654–69. Epub 2020/07/24. doi: 10.1039/d0ay00072h. PubMed PMID: 32701099.

28. Stella S, Vitale SR, Massimino M, Puma A, Tomarchio C, Pennisi MS, Tirro E, Romano C, Martorana F, Stagno F, Di Raimondo F, Manzella L. A Novel System for Semiautomatic Sample Processing in Chronic Myeloid Leukaemia: Increasing Throughput without Impacting on Molecular Monitoring at Time of SARSCoV-2 Pandemic. Diagnostics (Basel). 2021;11(8). Epub 2021/08/28. doi: 10.3390/diagnostics11081502. PubMed PMID: 34441436; PMCID: PMC8391152.

29. Dawes JC, Webster P, Iadarola B, Garcia-Diaz C, Dore M, Bolt BJ, Dewchand H, Dharmalingam G, McLatchie AP, Kaczor J, Caceres JJ, Paccanaro A, Game L, Parrinello S, Uren AG. LUMI-PCR: an Illumina platform ligation-mediated PCR protocol for integration site cloning, provides molecular quantitation of integration sites. Mob DNA. 2020;11:7. Epub 2020/02/12. doi: 10.1186/s13100-020-0201-4. PubMed PMID: 32042315; PMCID: PMC7001329.

30. Yu C, Caothien R, Jackson M, Nakao B, Pham A, Tam L, Roose-Girma M. Advanced Technologies and Automation in mES Cell Workflow. Methods Mol Biol. 2023;2631:183–206. Epub 2023/03/31. doi: 10.1007/978-1-0716-2990-1_7. PubMed PMID: 36995668.

31. Mardis E, McCombie WR. Preparing Polymerase Chain Reaction (PCR) Products for Capillary Sequencing. Cold Spring Harb Protoc. 2017;2017(7):pdb prot094599. Epub 2016/11/03. doi: 10.1101/pdb.prot094599. PubMed PMID: 27803282.

32. Santacruz D, Enane FO, Fundel-Clemens K, Giner M, Wolf G, Onstein S, Klimek C, Smith Z, Wijayawardena B, Viollet C. Automation of high-throughput mRNA-seq library preparation: a robust, hands-free and time efficient methodology. SLAS Discov. 2022;27(2):140–7. Epub 2022/01/31. doi: 10.1016/j.slasd.2022.01.002. PubMed PMID: 35093290.

33. Kind D, Baskaran P, Ramirez F, Giner M, Hayes M, Santacruz D, Koss CK, El Kasmi KC, Wijayawardena B, Viollet C. Automation enables high-throughput and reproducible single-cell transcriptomics library preparation. SLAS Technol. 2022;27(2):135–42. Epub 2022/01/22. doi: 10.1016/j.slast.2021.10.018. PubMed PMID: 35058211.

34. Arrigoni L, Ferrari F, Weller J, Bella C, Bonisch U, Manke T. AutoRELACS: automated generation and analysis of ultra-parallel ChIP-seq. Sci Rep. 2020;10(1):12400. Epub 2020/07/28. doi: 10.1038/s41598-020-69443-8. PubMed PMID: 32709929; PMCID: PMC7381599.

35. Malinowska JM, Palosaari T, Sund J, Carpi D, Lloyd GR, Weber RJM, Whelan M, Viant MR. Automated Sample Preparation and Data Collection Workflow for High- Throughput In Vitro Metabolomics. Metabolites. 2022;12(1). Epub 2022/01/21. doi: 10.3390/metabo12010052. PubMed PMID: 35050173; PMCID: PMC8778710.

36. Chen Y, Kaplan Lease N, Gin JW, Ogorzalek TL, Adams PD, Hillson NJ, Petzold CJ. Modular automated bottom-up proteomic sample preparation for high-throughput applications. PLoS One. 2022;17(2):e0264467. Epub 2022/02/26. doi: 10.1371/journal.pone.0264467. PubMed PMID: 35213656; PMCID: PMC8880914.

37. Pajak L, Zhang R, Pittman C, Roby K, Boyer S. Automated Genomic and Proteomic Applications on the Biomek® NX Laboratory Automation Workstation. SLAS Technology. 2004;9(3):177–84. doi: 10.1016/j.jala.2004.04.006.

38. Wu Q, Sui X, Tian R. [Advances in high-throughput proteomic analysis]. Se Pu. 2021;39(2):112–7. Epub 2021/07/07. doi: 10.3724/SP.J.1123.2020.08023. PubMed PMID: 34227342; PMCID: PMC9274848

39. Zhu M, Zhang P, Geng-Spyropoulos M, Moaddel R, Semba RD, Ferrucci L. A robotic protocol for highthroughput processing of samples for selected reaction monitoring assays. Proteomics. 2017;17(6). Epub 2016/11/20. doi: 10.1002/pmic.201600339. PubMed PMID: 27862927; PMCID: PMC5534325.

40. Ball M, Romanovsky E, Schnecko F, Kirchner M, Neumann O, Brandt R, Beck S, Seker-Cin H, Kluck K, Ourailidis I, Goldschmid H, Fink A, Volckmar AL, Menzel M, Allgäuer M, Schirmacher P, Budczies J, Stenzinger A, Kazdal D. Clinical Implementation of a High-Throughput Automated Comprehensive Genomic Profiling Test: TruSight Oncology 500 HT. J Mol Diagn. 2025 Feb;27(2):154–162. doi: 10.1016/j.jmoldx.2024.11.005. Epub 2024 Dec 12. PMID: 39674366; PMCID: PMC12179514.

41. Uhlén M, Fagerberg L, Hallström BM, Lindskog C, Oksvold P, Mardinoglu A, Sivertsson Å, Kampf C, Sjöstedt E, Asplund A, Olsson I, Edlund K, Lundberg E, Navani S, Szigyarto CA, Odeberg J, Djureinovic D, Takanen JO, Hober S, Alm T, Edqvist PH, Berling H, Tegel H, Mulder J, Rockberg J, Nilsson P, Schwenk JM, Hamsten M, von Feilitzen K, Forsberg M, Persson L, Johansson F, Zwahlen M, von Heijne G, Nielsen J, Pontén F. Proteomics. Tissue-based map of the human proteome. Science. 2015 Jan 23;347(6220):1260419. doi: 10.1126/science.1260419. PMID: 25613900.

42. Jaluria P, Betenbaugh M, Konstantopoulos K, Shiloach J. Enhancement of cell proliferation in various mammalian cell lines by gene insertion of a cyclin-dependent kinase homolog. BMC Biotechnol. 2007 Oct 18;7:71. doi: 10.1186/1472-6750-7-71. PMID: 17945021; PMCID: PMC2164945.

43. Drake JM, Graham NA, Stoyanova T, Sedghi A, Goldstein AS, Cai H, Smith DA, Zhang H, Komisopoulou E, Huang J, Graeber TG, Witte ON. Oncogene-specific activation of tyrosine kinase networks during prostate cancer progression. Proc Natl Acad Sci U S A. 2012 Jan 31;109(5):1643–8. doi: 10.1073/pnas.1120985109. Epub 2012 Jan 17. PMID: 22307624; PMCID: PMC3277127.

44. Haneke K, Schott J, Lindner D, Hollensen AK, Damgaard CK, Mongis C, Knop M, Palm W, Ruggieri A, Stoecklin G. CDK1 couples proliferation with protein synthesis. J Cell Biol. 2020 Mar 2;219(3):e201906147. doi: 10.1083/jcb.201906147. PMID: 32040547; PMCID: PMC7054999.

45. Siddika T, Balasuriya N, Frederick MI, Rozik P, Heinemann IU, O’Donoghue P. Delivery of Active AKT1 to Human Cells. Cells. 2022 Nov 29;11(23):3834. doi: 10.3390/cells11233834. PMID: 36497091; PMCID: PMC9738475.

46. Upadhya D, Ogata M, Reneker LW. MAPK1 is required for establishing the pattern of cell proliferation and for cell survival during lens development. Development. 2013 Apr;140(7):1573–82. doi: 10.1242/dev.081042. PMID: 23482492; PMCID: PMC3596996.

47. Greulich H, Erikson RL. An analysis of Mek1 signaling in cell proliferation and transformation. J Biol Chem. 1998 May 22;273(21):13280–8. doi: 10.1074/jbc.273.21.13280. PMID: 9582373.

48. Koff A, Giordano A, Desai D, Yamashita K, Harper JW, Elledge S, Nishimoto T, Morgan DO, Franza BR, Roberts JM. Formation and activation of a cyclin E-cdk2 complex during the G1 phase of the human cell cycle. Science. 1992 Sep 18;257(5077):1689–94. doi: 10.1126/science.1388288. PMID: 1388288.

49. Gheghiani L, Loew D, Lombard B, Mansfeld J, Gavet O. PLK1 Activation in Late G2 Sets Up Commitment to Mitosis. Cell Rep. 2017 Jun 6;19(10):2060–2073. doi: 10.1016/j.celrep.2017.05.031. PMID: 28591578.

50. Jeong AL, Han S, Lee S, Su Park J, Lu Y, Yu S, Li J, Chun KH, Mills GB, Yang Y. Patient derived mutation W257G of PPP2R1A enhances cancer cell migration through SRC-JNK-c-Jun pathway. Sci Rep. 2016 Jun 7;6:27391. doi: 10.1038/srep27391. PMID: 27272709; PMCID: PMC4895347.

51. Bhardwaj A, Singh S, Srivastava SK, Arora S, Hyde SJ, Andrews J, Grizzle WE, Singh AP. Restoration of PPP2CA expression reverses epithelial-to-mesenchymal transition and suppresses prostate tumour growth and metastasis in an orthotopic mouse model. Br J Cancer. 2014 Apr 15;110(8):2000–10. doi: 10.1038/bjc.2014.141. Epub 2014 Mar 18. PMID: 24642616; PMCID: PMC3992501.

52. Yong L, YuFeng Z, Guang B. Association between PPP2CA expression and colorectal cancer prognosis tumor marker prognostic study. Int J Surg. 2018 Nov;59:80–89. doi: 10.1016/j.ijsu.2018.09.020. Epub 2018 Oct 5. PMID: 30296597.

53. Xiao T, Xiao Y, Wang W, Tang YY, Xiao Z, Su M. Targeting EPHA2 in cancer. J Hematol Oncol. 2020 Aug 18;13(1):114. doi: 10.1186/s13045-020-00944-9. PMID: 32811512; PMCID: PMC7433191.

54. Wilson K, Shiuan E, Brantley-Sieders DM. Oncogenic functions and therapeutic targeting of EPHA2 in cancer. Oncogene. 2021 Apr;40(14):2483–2495. doi: 10.1038/s41388-021-01714-8. Epub 2021 Mar 8. PMID: 33686241; PMCID: PMC8035212.

55. Zelinski DP, Zantek ND, Stewart JC, Irizarry AR, Kinch MS. EPHA2 overexpression causes tumorigenesis of mammary epithelial cells. Cancer Res. 2001 Mar 1;61(5):2301–6. PMID: 11280802.

56. Park JE, Son AI, Zhou R. Roles of EPHA2 in Development and Disease. Genes (Basel). 2013 Jul 1;4(3):334–57. doi: 10.3390/genes4030334. PMID: 24705208; PMCID: PMC3924825.

57. Wilson K, Shiuan E, Brantley-Sieders DM. Oncogenic functions and therapeutic targeting of EPHA2 in cancer. Oncogene. 2021 Apr;40(14):2483–2495. doi: 10.1038/s41388-021-01714-8. Epub 2021 Mar 8. PMID: 33686241; PMCID: PMC8035212.

58. Browne CD, Hoefer MM, Chintalapati SK, Cato MH, Wallez Y, Ostertag DV, Pasquale EB, Rickert RC. SHEP1 partners with CasL to promote marginal zone B-cell maturation. Proc Natl Acad Sci U S A. 2010 Nov 2;107(44):18944–9. doi: 10.1073/pnas.1007558107. Epub 2010 Oct 18. PMID: 20956287; PMCID: PMC2973925.

59. Ireton RC, Chen J. EPHA2 receptor tyrosine kinase as a promising target for cancer therapeutics. Curr Cancer Drug Targets. 2005 May;5(3):149–57. doi: 10.2174/1568009053765780. PMID: 15892616.

60. Zhang G, Njauw CN, Park JM, Naruse C, Asano M, Tsao H. EPHA2 is an essential mediator of UV radiation-induced apoptosis. Cancer Res. 2008 Mar 15;68(6):1691–6. doi: 10.1158/0008-5472.CAN-07-2372. PMID: 18339848; PMCID: PMC4469360.

61. Pasquale EB. Eph receptors and ephrins in cancer progression. Nat Rev Cancer. 2024 Jan;24(1):5–27. doi: 10.1038/s41568-023-00634-x. Epub 2023 Nov 23. PMID: 37996538; PMCID: PMC11015936.

62. Miao H, Wei BR, Peehl DM, Li Q, Alexandrou T, Schelling JR, Rhim JS, Sedor JR, Burnett E, Wang B. Activation of EphA receptor tyrosine kinase inhibits the Ras/MAPK pathway. Nat Cell Biol. 2001 May;3(5):527–30. doi: 10.1038/35074604. PMID: 11331884.

63. Noberini R, Mitra S, Salvucci O, Valencia F, Duggineni S, Prigozhina N, Wei K, Tosato G, Huang Z, Pasquale EB. PEGylation potentiates the effectiveness of an antagonistic peptide that targets the EphB4 receptor with nanomolar affinity. PLoS One. 2011;6(12):e28611. doi: 10.1371/journal.pone.0028611. Epub 2011 Dec 14. PMID: 22194865; PMCID: PMC3237458.

64. Pasquale EB. Eph receptor signalling casts a wide net on cell behaviour. Nat Rev Mol Cell Biol. 2005 Jun;6(6):462–75. doi: 10.1038/nrm1662. Erratum in: Nat Rev Mol Cell Biol. 2005 Jul;6(7):589. PMID: 15928710.

65. Lin S, Gordon K, Kaplan N, Getsios S. Ligand targeting of EphA2 enhances keratinocyte adhesion and differentiation via desmoglein 1. Mol Biol Cell. 2010 Nov 15;21(22):3902–14. doi: 10.1091/mbc.E10-03-0242. Epub 2010 Sep 22. PMID: 20861311; PMCID: PMC2982116.

66. Singh DR, Ahmed F, King C, Gupta N, Salotto M, Pasquale EB, Hristova K. EphA2 Receptor Unliganded Dimers Suppress EphA2 Pro-tumorigenic Signaling. J Biol Chem. 2015 Nov 6;290(45):27271–27279. doi: 10.1074/jbc.M115.676866. Epub 2015 Sep 11. PMID: 26363067; PMCID: PMC4646390.

67. Singh DR, Ahmed F, Paul MD, Gedam M, Pasquale EB, Hristova K. The SAM domain inhibits EphA2 interactions in the plasma membrane. Biochim Biophys Acta Mol Cell Res. 2017 Jan;1864(1):31–38. doi: 10.1016/j.bbamcr.2016.10.011. Epub 2016 Oct 21. PMID: 27776928; PMCID: PMC5148718.

68. Miura K, Nam JM, Kojima C, Mochizuki N, Sabe H. EphA2 engages Git1 to suppress Arf6 activity modulating epithelial cell-cell contacts. Mol Biol Cell. 2009 Apr;20(7):1949–59. doi: 10.1091/mbc.e08-06-0549. Epub 2009 Feb 4. PMID: 19193766; PMCID: PMC2663931.

69. Zhang G, Njauw CN, Park JM, Naruse C, Asano M, Tsao H. EphA2 is an Essential Mediator of Ultraviolet Radiation-induced Apoptosis. Cancer research (Chicago, Ill.), 2008-03, Vol.68 (6), p.1691-1696. DOI: 10.1158/0008-5472.CAN-07-2372. PMID: 18339848.

