## Supplementary for "Automation of high-throughput workflow for arrayed CRISPR activation library screening"

**Supplementary Table 1. PCR primer sequences.**

| | |
|---|---|
| **EphA2_F** | AGAGGCTGAGCGTATCTTCAT |
| **EphA2_R** | GGTCCGACTCGGCATAGTAGA |
| **POMK_F** | CTTCTTCATCGCTCCTCGACA |
| **POMK_R** | AGCCAAGGTGAGCAGTTTTTC |
| **CSNK1A1L_F** | CTTCTTGTCTGTAAGCCAGC |
| **CSNK1A1L_R** | TCTTATGTCTTCACAGGTAAGC |
| **ROCK1_F** | CAAGGTGGTGATGGTTATTATG |
| **ROCK1_R** | TATCACCTACAAGCATTTCG |
| **PTPRO_F** | ATGACTTCAGCCGTGTGAGA |
| **PTPRO_R** | TGTTGCAGGACCATCTTCCA |
| **GSK3B_F** | GGCAGCATGAAAGTTAGCAGA |
| **GSK3B_R** | GGCGACCAGTTCTCCTGAATC |
| **RIOK2_F** | ACAACAGGCAAGATGGTCA |
| **RIOK2_R** | GACGACAAGGCAATTAGATGAG |
| **MARK3_F** | CGCTCTCTGCTCCTCCTGTT |
| **MARK3_R** | TGCTGGTCTGACTCCTTTTCGG |
| **HPRT_F*** | TGGAGTCCTATTGACATCGCCAGT |
| **HPRT_R*** | AACAACAATCCGCCCAAAGGGAAC |

* HPRT serves as a housekeeping gene for normalization control.

**Supplementary Table 2. Guide-RNA sequences.**

| | |
|---|---|
| **EphA2-qgRNAs** | gCCCCTACGGATTAGCCCCC<br>gGGGAGCCCGCGAGTCCAGA<br>GGAATGTCTTTAAAGGGGCC<br>gGGCATGAATGAACAGGAGT |
| **EphA2-sgRNA-1** | gCCCCTACGGATTAGCCCCC |
| **EphA2-sgRNA-2** | gGGGAGCCCGCGAGTCCAGA |
| **EphA2-sgRNA-3** | GGAATGTCTTTAAAGGGGCC |
| **EphA2-sgRNA-4** | gGGCATGAATGAACAGGAGT |
| **NS6-qgRNAs** | TAGTCTCACCTGATGGCGTG<br>cCATAATATACGCCCGGCAT<br>tACCCAACGTCCGAGAGGGT<br>AGCTAGCGATGGCTCTAAGT |
| **NS10-qgRNAs** | GACGAGAGAAATTCAACGCA<br>aCCGCTGAGAGCCATATTTG<br>GTGGCACGGCCACGTAAAAC<br>TCTACGTGTAGTTGTACATA |
| **NS12-qgRNAs** | TTCGGAACTTACTCAGGGTA<br>ATTAGCCGTTGCCATATCAA<br>tTTGACCGTTATAGTCTTAT<br>AGGGCGAGCAGCAGAGTACG |

Quadruple-guide RNAs (gqRNAs); Single-guide RNA (sgRNA).
